# GCNCDA: A New Method for Predicting CircRNA-Disease Associations Based on Graph Convolutional Network Algorithm

**DOI:** 10.1101/858837

**Authors:** Lei Wang, Zhu-Hong You, Yang-Ming Li, Kai Zheng, Yu-An Huang

**Affiliations:** Xinjiang Technical Institutes of Physics and Chemistry, Chinese Academy of Sciences, Urumqi, 830011, China; Department of Electrical Computer and Telecommunications Engineering Technology, Rochester Institute of Technology, Rochester, 14623, USA; School of Computer Science and Technology, China University of Mining and Technology, Xuzhou, 221116, China; Department of Computing, Hong Kong Polytechnic University, Hong Kong 999077, China

## Abstract

Numerous evidences indicate that Circular RNAs (circRNAs) are widely involved in the occurrence and development of diseases. Identifying the association between circRNAs and diseases plays a crucial role in exploring the pathogenesis of complex diseases and improving the diagnosis and treatment of diseases. However, due to the complex mechanisms between circRNAs and diseases, it is expensive and time-consuming to discover the new circRNA-disease associations by biological experiment. Therefore, there is increasingly urgent need for utilizing the computational methods to predict novel circRNA-disease associations. In this study, we propose a computational method called GCNCDA based on the deep learning Fast learning with Graph Convolutional Networks (FastGCN) algorithm to predict the potential disease-associated circRNAs. Specifically, the method first forms the unified descriptor by fusing disease semantic similarity information, disease and circRNA Gaussian Interaction Profile (GIP) kernel similarity information based on known circRNA-disease associations. The FastGCN algorithm is then used to objectively extract the high-level features contained in the fusion descriptor. Finally, the new circRNA-disease associations are accurately predicted by the Forest by Penalizing Attributes (Forest PA) classifier. The 5-fold cross-validation experiment of GCNCDA achieved 91.2% accuracy with 92.78% sensitivity at the AUC of 90.90% on circR2Disease benchmark dataset. In comparison with different classifier models, feature extraction models and other state-of-the-art methods, GCNCDA shows strong competitiveness. Furthermore, 10 of the top 15 circRNA-disease association candidates with the highest prediction scores were confirmed by recently published literature. These results suggest that GCNCDA can effectively predict potential circRNA-disease associations and provide highly credible candidates for biological experiments.

**Author Summary:** The recognition of circRNA-disease association is the key of disease diagnosis and treatment, and it is of great significance for exploring the pathogenesis of complex diseases. Computational methods can predicte the potential disease-related circRNAs quickly and accurately. Based on the hypothesis that circRNA with similar function tends to associate with similar disease, GCNCDA model is proposed to effectively predict the potential association between circRNAs and diseases by combining FastGCN algorithm. The performance of the model was verified by cross-validation experiments, different feature extraction algorithm and classifier models comparison experiments. Furthermore, 10 of the top 15 disease-associated circRNAs with the highest prediction scores were confirmed by recently published literature. It is anticipated that GCNCDA model can give priority to the most promising circRNA-disease associations on a large scale to provide reliable candidates for further biological experiment.

## 1. Introduction

As a new type of endogenous non-coding RNA, circular RNA (circRNA) has a closed-loop structure without a 5’and 3’polyadenylated tails [1-3]. As early as 1971, researchers discovered the viroids genome composed of single-stranded closed RNA molecules in potatoes [4]. In 1979, Hsu *et al.* [5] observed the presence of circRNA in the cytoplasm of eukaryotic cells by electron microscopy. In 1995, the researchers [6] found that the mouse sperm determinant gene Sry has circular transcription during transcription. But these findings did not attract much attention of researchers at the time. Until 2012, Salzman *et al.* [7] reported about 80 circRNAs for the first time with the help of high-throughput sequencing technology. Since then, a large number of circRNA molecules have been identified.

With the rapid development of bioinformatics and the continuous innovation of high-throughput sequencing technology, a large number of endogenous circRNA have been found in eukaryotic cells. CircRNA has the characteristics of universality, conservativeness, tissue-specificity and stability. Its unique sequence structure makes it have the functions of microRNA sponge [8], regulators of RNA binding proteins[9] and transcription of parental genes [10]. In addition, it is involved in the development and progression of diseases such as cancer [11,12], diabetes [13], nervous system diseases [14] and atherosclerosis [15]. For example, Burd *et al.* [16] found that the expression of cANRIL (circular antisense non-coding RNA in the INK4 locus) is an antisense transcript of INK4/ARF gene, which can inhibit the expression of INK4/ARF through specific multi comb family complex, thereby affecting the risk of atherosclerosis. Du *et al.* [17] found that circ-Foxo3, a member of the transcription factor foxo3, is highly expressed in myocardial samples from elderly patients and rats. It can prevent and reposition ID-1, E2F1, FAK and H1F1a in the cytoplasm and prevent their anti-aging function. By establishing the HT22 cell model of oxygen-glucose deprivation/reoxygenation (OGD/R), Lin *et al.* [18] found that the expression of mmu-circRNA-015947 was higher than that of normal cells, indicating that the expression of circRNA was involved in OGD/R-induced neuron injury. Lukiw [19] found that in the hippocampal CA1 region of Alzheimer’s disease (AD), there is a dysregulation of the miRNA-circRNA system. When the expression of CDRlas (CiRS-7) decreased or the ability to adsorb microRNA-7 weakened, the expression of miR-7 is increased and directly leads to down-regulation of ubiquitin ligase an expression in the human central nervous system, thereby affecting the normal function of the central nervous system and causing serious damage to brain tissue. Numerous studies have shown that circRNA can be a new clinical diagnostic marker or a potential target for human disease treatment. Therefore, the identification of disease-related circRNA may help to reveal the mechanism of disease occurrence and development, and further promote the understanding of complex human diseases.

As the number of detected circRNAs increases, multiple databases have been created to store information on circRNAs, such as Circ2Traits [20], circBase [21], deepBase [22] and CircNet [23]. Furthermore, researchers have gradually collected circRNA-disease associations supported by experiments and established databases, such as circR2Disease [24], circRNADb [25], circRNADisease [26] and Circ2Disease [27]. The accumulation of these data provides an opportunity for computational methods to predict potential circRNA-disease associations. For example, Xiao *et al.* [28] proposed an integrated computational framework called MRLDC to identify disease-associated circRNAs based on the hypothesis that circRNAs with similar functions are usually associated with similar diseases, and vice versa. Yan *et al.* [29] developed the DWNN-RLS method using Regularized Least Squares of Kronecker product kernel to predict circRNA-disease associations. In the experiment, this method achieved AUC of 0.8854, 0.9205 and 0.9701 in 5-fold CV, 10-fold CV and LOOCV, respectively. Fan *et al.* [30] proposed the KATZHCDA model for predicting circRNA-disease associations based on a heterogeneous network constructed by disease phenotype similarity, circRNA expression profiles and Gaussian interaction profile kernel similarity. As a result, KATZHCDA reached the AUC values of 0.7936 and 0.8469 in 5-fold cross-validation and LOOCV, respectively. Although the above models play important roles in the development of circRNA-disease association prediction computational methods and have achieved fruitful results, they are limited by certain problems: (1) the existing data are derived from incompletely related biological information, which cannot fully describe the complex association between circRNA and disease. (2) The experimentally verified circRNA-disease associations are limited in number and have some noise information, which easily leads to many false negative associations predicted by the model.

The purpose of this study is to propose a new computational model to predict the potential circRNA-disease associations in an attempt to overcome these problems. The proposed model GCNCDA has the following advantages: (1) Comprehensive use of disease semantic similarity information, disease GIP kernel similarity information, circRNA GIP kernel similarity information and known circRNA-disease association information to accurately predict potential circRNA-disease associations. (2) The advanced features of circRNA-disease associations are extracted by the deep learning FastGCN algorithm to reduce false negative associations and improve model performance. In the 5-fold cross-validation experiment on the benchmark dataset, GCNCDA achieved an AUC value of 90.90%. The results of comparative experiments show that GCNCDA is superior to other competing models and can effectively predict potential circRNA-disease associations. Furthermore, case studies show that GCNCDA can identify new circRNA-disease associations, which are validated by the latest literature. It is worth noting that the performance of GCNCDA is underestimated due to experimentally verified limitations on the number of circRNA-disease associations.

## 2. Materials and Methods

### 2.1 Method overview

In this study, we propose a computational method called GCNCDA to predict potential circRNA-disease associations. The execution process of GCNCDA is divided into three steps, and its framework is shown in Figure 1. Firstly, we construct descriptor by synthesizing disease semantic similarity information, disease GIP kernel similarity information, circRNA GIP kernel similarity information and known circRNA-disease association information. Secondly, the FastGCN algorithm of deep learning is used to extract high-level features of descriptors and reduce noise information. Finally, the extracted high-level features are fed into Forest PA classifier to accurately predict the potential circRNA-disease associations.

**Figure 1.**
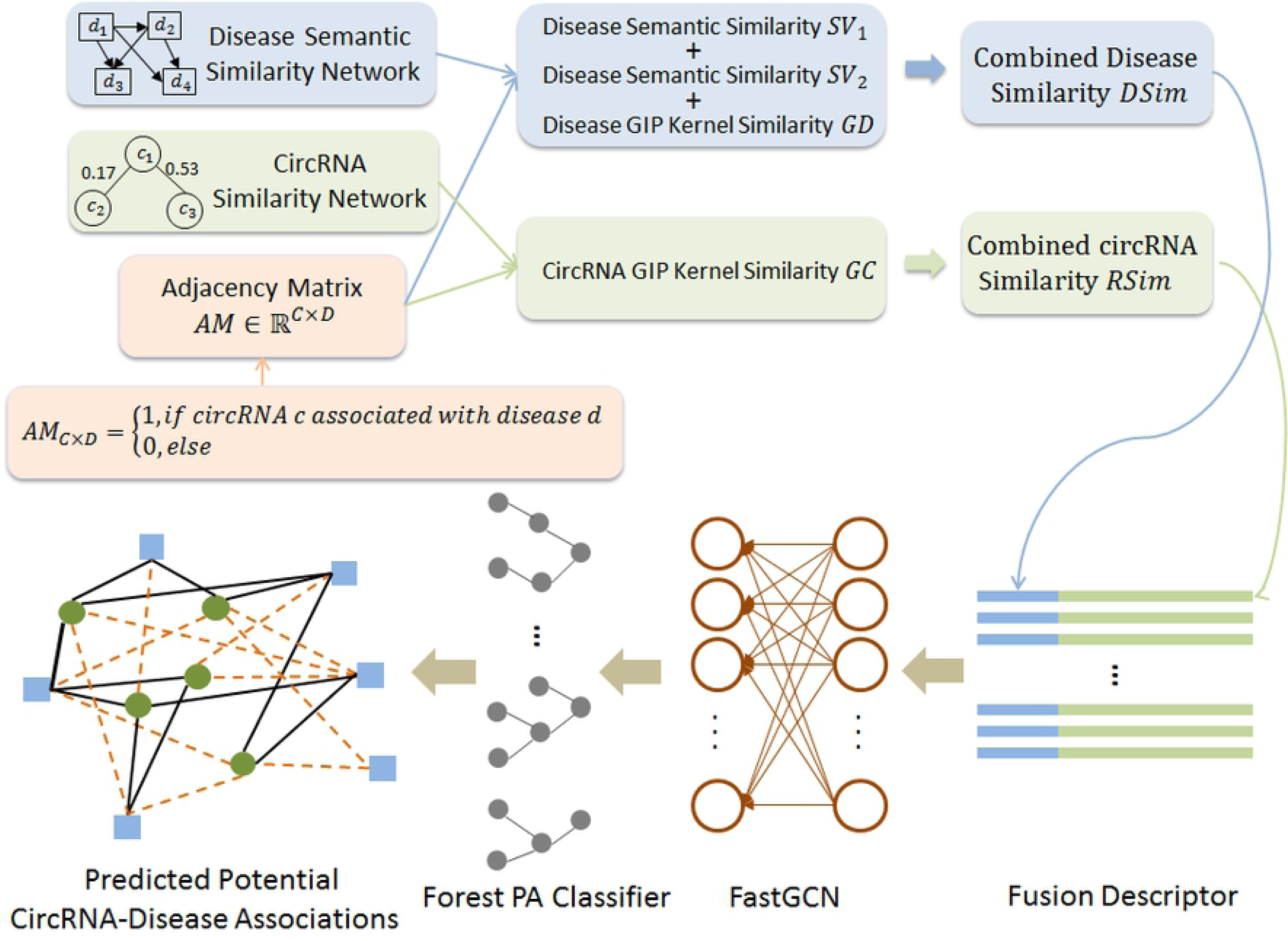
The framework of GCNCDA to predict potential circRNA-disease associations

### 2.2 Benchmark Dataset

In this study, we used the recently established experimentally verified circRNA-disease association dataset circR2Disease [24] as the benchmark dataset to evaluate the performance of various models. CircR2Disease is a dedicated database and comprehensive platform that collects disease-related circRNAs from experimental support. The database currently hosts 739 entries from published literature, including 661 circRNAs, and 100 diseases. The benchmark dataset can be expressed as:

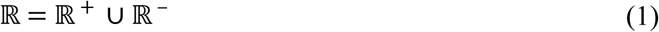

where ∪ denotes the union symbol in set theory, ℝ^+^ represents the positive dataset, which contains 739 circRNA-disease associations with experimentally verified, ℝ^−^ represents the negative dataset, which contains 739 circRNA-disease associations without experimentally verified. The circR2Disease dataset can be available on the website http://bioinfo.snnu.edu.cn/CircR2Disease/.

In the circR2Disease dataset, there were a total of 661 × 100 − 739 = 65361 circRNA-disease associations without experimental verified. If they are all treated as negative samples, they will form an unbalanced dataset. In order to avoid bias in the prediction results caused by unbalanced data, we solve this problem by reducing the number of negative samples by the down-sampling method. Specifically, we select 739 negative samples from all negative samples using random sampling without replacement, and then combine the positive samples to form a distributed equilibrium dataset. In theory, there may be unconfirmed circRNA-disease associations in these 65361 negative samples. But in the 739 negative samples we selected, this probability is much less than 739 ÷ (661 × 100 − 739) ≈ 1.13%. Thus, we constructed the dataset containing 1478 samples in this way, in which the number of positive samples is the same as that of negative samples. Known circRNA-disease associations and their names obtatined from circR2Disease database can be seen in Supplementary Tables 1-3.

Based on the circR2Disease dataset, we constructed 661 × 100 dimensional adjacency matrix *AM*, where 661 represents the number of circRNAs, and 100 represents the number of diseases. When circRNA *c*(*i*) is associated with disease *d*(*j*), element *AM*(*i,j*) of matrix *AM* is assigned a value of 1. Otherwise, it is assigned a value of 0.

### 2.3 Construction of CircRNA Similarity Model

In this study, we used the Gaussian interaction profile (GIP) kernel similarity to construct the similarity model of circRNA. Based on the hypothesis that circRNAs with similar function are often associated with similar diseases, and vice versa, we established the GIP kernel similarity model of circRNA according to the known circRNA-disease association network. Specifically, we define the binary vector *V*(*c*(*i*)) to represent the interaction profiles of circRNA *c*(*i*). The dimension of the vector *V*(c(*i*)) is 100, which corresponds to 100 diseases in adjacent matrix *AM*. When disease *c*(*i*) is associated with one of 100 diseases, the corresponding bit in vector *V*(*c*(*i*)) is set to 1. Otherwise, it is set to 0. That is to say, the interaction profiles binary vector *V*(*c*(*i*)) is the row vector of the row corresponding to disease c(*i*) in the adjacency matrix *AM*. Thus, we can get the circRNA GIP kernel similarity *GC*(*c*(*i*),*c*(*j*)) of circRNA *c*(*i*) and circRNA *c*(*j*):

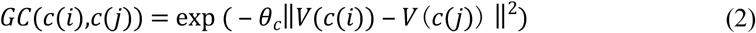

where *θ*_*c*_ is the width parameter, which can be calculated using the normalized original parameters of the following formula:

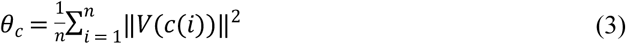

where *n* is the column number of adjacent matrix *AM*.

### 2.4 Construction of Disease Similarity Model

The disease similarity model consists of two parts: the disease GIP kernel similarity and the disease semantic similarity. For the disease GIP kernel similarity, our construction method is similar to the GIP kernel similarity of circRNA. More concretely, we define a binary vector *V*(*d*(*i*)) to represent the interaction profiles of disease *d*(*i*) according to the adjacent matrix *AM* provided by circR2Disease dataset. The dimension of the vector *V*(*d*(*i*)) is 661, which corresponds to 661 circRNAs in adjacent matrix *AM*. When disease *d*(*i*) is associated with one of 661 circRNAs, the corresponding bit in vector *V*(*d*(*i*)) is set to 1. Otherwise, it is set to 0. That is to say, the interaction profiles binary vector *V*(*d*(*i*)) is the column vector of the column corresponding to disease *d*(*i*) in the adjacency matrix *AM*. Through the above definition, we can calculate the disease GIP kernel similarity *GD*(*d*(*i*),*d*(*j*)) of disease *d*(*i*) and disease *d*(*j*):

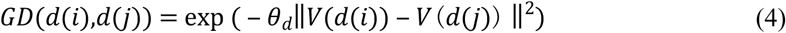

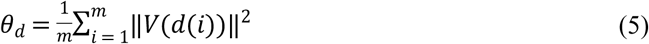

where *θ*_*d*_ is the width parameter and *m* is the row number of adjacent matrix *AM*

For disease semantic similarity, we construct it through the MeSH database [31-33] from the National Library of Medicine (NLM). It can be downloaded at https://www.nlm.nih.gov/. The MeSH database gives a rigorous disease classification system that uses a Directed Acyclic Graph (DAG) to reflect relationships between different diseases. The MeSH dataset can be seen in Supplementary Table 4. In DAG, a node represents disease, and an edge represents the relationship between diseases. Given a disease *d* whose structure can be expressed as *DAG*_*d*_ = (*d,N*_*d*_,*E*_*d*_), where *N*_*d*_ represents the set of diseases associated with *d* including disease *d* itself, and *E*_*d*_ represents the relationship between these diseases. For a disease *s* within *DAG*_*d*_, its contribution value *D*_*d*_(*s*) can be calculated by the following formula:

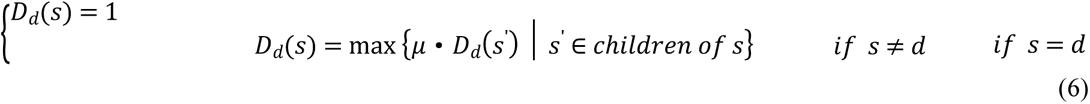

where *μ* indicates the semantic contribution factor between disease *s* and its child disease *s*’. According to the previous study by Wang *et al.* [34], we set the semantic contribution factor *μ* to the optimal value of 0.5. Thus, by accumulating the contribution values of all children with disease *d*, we can get their semantic values *DV*(*d*):

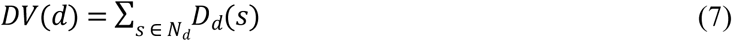

In general, the more nodes that are shared between DAGs of different diseases, the more similar they are. Based on this assumption, we construct the first disease semantic similarity model *SV*_1_(*d*(*i*),*d*(*j*)) of disease *d*(*i*) and disease *d*(*j*) through the DAG hierarchical relationship of disease:

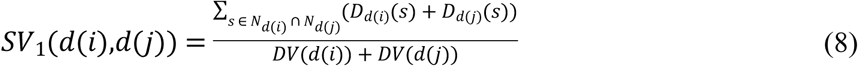

In disease semantic similarity model *SV*_1_, we mainly consider the hierarchical relationship of disease DAG, that is, the disease in the same layer in the DAG contributes the same value to the disease *d*. However, the number of different diseases in DAGs can also affect the semantic similarity of disease. The fewer diseases appear in DAGs, the more important they are. Therefore, we constructed the second method for calculating the disease contribution value based on this hypothesis:

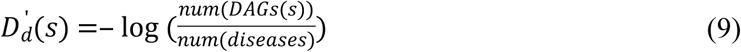

where *num*(*DAGs*(*s*)) denotes the number of DAGs that contain disease *s*, and *num*(*diseases*) denotes the number of all diseases. Thus, the second disease semantic similarity model *SV*_2_(*d*(*i*),*d*(*j*)) of disease *d*(*i*) and disease *d*(*j*) can be calculated as follows:

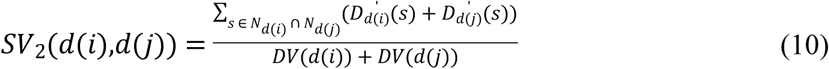

where *DV*(*d*(*i*)) and *DV*(*d*(*j*)) have the same meaning as disease semantic similarity model *SV*_1_, which can be calculated from formula 7.

### 2.5 Multi-source Data Fusion

In order to make full use of information from different sources, we used the fusion method to fuse circRNA similarity information and disease similarity information with known circRNA-disease associations. The fused information can absorb the characteristics of different data sources, thus describing the complex relationship between circRNAs and diseases more comprehensively.

For the circRNA, we use the constructed circRNA GIP kernel similarity *GR* directly to represent the circRNA descriptor *RSim*. For the disease, we need to fuse the disease semantic similarity model *SV*_1_ and *SV*_2_, and disease GIP kernel similarity *GD*. Since the MeSH database provides a strict disease association, we use it as much as possible. More specifically, if there is the semantic similarity between disease *d*(*i*) and disease *d*(*j*), then the disease semantic similarity is used to construct the descriptor *DSim*. Otherwise, it is constructed using disease GIP kernel similarity. This construction rule can be described by the following formula:

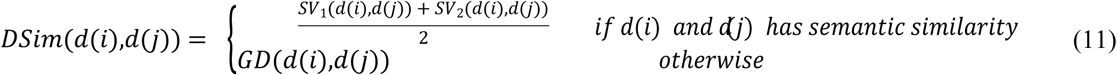

Finally, we match circRNA similarity *RSim* with disease similarity *DSim* based on known circRNA-disease associations to form a complete fusion descriptor. The fusion descriptor FV (*c*(*i*),*d*(*j*)) of circRNA *c*(*i*) and disease *d*(*j*) can be described as follows:

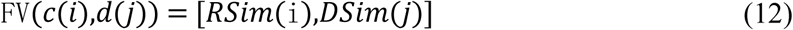

where *RSim*(*i*) indicates the i row vector of circRNA *c*(*i*) in the circRNA similarity matrix *RSim*, and *DSim*(*j*) indicates the *j* column vector of disease *d*(*j*) in the disease similarity matrix *DSim*.

### 2.6 Feature Extraction by Fast Learning with Graph Convolutional Networks

After getting the fusion descriptors, we used the Fast learning with Graph Convolutional Networks (FastGCN) [35] algorithm to extract their features to remove noise information and improve the performance of the model. FastGCN is an efficient algorithm based on the original GCN and realized by importance sampling. It interprets graph convolutions as integral transforms of embedding functions under probability measure. To be specific, FastGCN interprets the graph vertices as independent and identically distributed (i.i.d.) samples of some probability distributions, and integrates loss and each convolution layer as vertex embedding functions. The integrals are then calculated by Monte Carlo approximation to determine the sample loss and sample gradient. Finally, important sampling is used to reduce the approximate variance. FastGCN not only eliminates the reliance on test data but also produces a controllable cost for each batch of computation.

Suppose there is a graph *G’* with the vertex set *V’* associated with a probability space (*V’, F,P*). For the given graph *G*, it is a subgraph of *G’* whose vertices are i.i.d. samples of *V’* obtained from the probability measure *P*. For the probability space, *V’* is used as the sample space, and *F* can be any event space. The probability measure *P* defines a sample distribution. Thus, the function generalization can be expressed as:

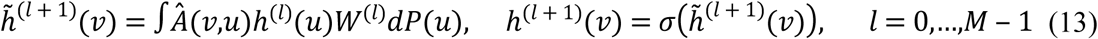

where the function *h*^(*l*)^ represents an embedding function from the *lth* layer, *u* and *v* are independent random variables that have the same probability measure *P*. The embedding functions of two consecutive layers are correlated by convolution and expressed by an integral transforma, where the kernel *Â*(*v,u*) corresponds to the (*v,u*) element of the matrix *Â*. The loss *L* is the expected value of g(*h*^(*M*)^) that is finally embedded in *h*^(*M*)^, and can be expressed as:

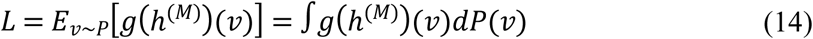

For the lth layer, the t_l_ i.i.d. sample 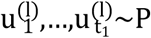 is used to approximatively estimate the integral transformation:

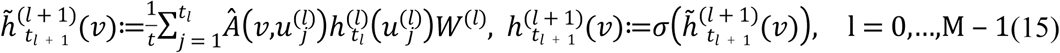

Here, 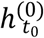 is *h*^(0)^. Therefore, the loss *L* is transformed into:

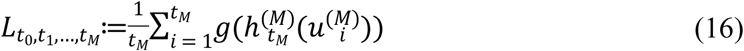

### 2.7 Prediction by Forest PA Classifier

In the experiment, we send the extracted features into the Forest by Penalizing Attributes (Forest PA) classifier for classification, so as to obtain accurate circRNA-disease association prediction results. Forest PA is a novel decision forest building algorithm recently proposed by Adnan *et al.* [36]. The Forest PA algorithm uses the complete attribute set to generate decision trees by imposing penalties on attributes participating in the latest decision tree. Besides, the participating attributes obtain random weights from the range of weights associated with the respective levels in the tree, thereby maintaining the decision tree generated by the algorithm with individually accuracy and diversity. The execution steps of the Forest PA algorithm are as follows:

1. The Forest PA first generates a bootstrap sample *D*_*i*_ from the original training data set *D*.
2. The Forest PA then uses the weight of attributes to generate decision trees from the bootstrap sample. When choosing the splitting attributes, Forest PA uses the CART algorithm with merit values, whose value is obtained by multiplying its classification ability with its weight.
3. The incremental values of attribute weights and gradient weight in the latest tree are updated iteratively. Here, the weights of the attributes appear in the latest tree will be updated. The weights of attributes that do not appear in the latest tree remain unchanged. Considering that the weight of attribute is determined by the level *λ* of test attributes in the latest tree, if an attribute appears on the root node, their value of *λ* is 1; if an attribute appears on the child node, their value of *λ* is 2. According to the value of *λ*, the weight of randomly generated attributes within a Weight-Range *WR* is defined as follows:

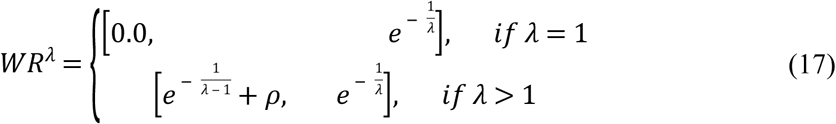
4. Update weights of the applicable attributes with the corresponding weight increment values that do not exist in the latest tree.

## 3. Results and discussion

### 3.1 Evaluation Criteria

In this study, we used the 5-fold cross-validation (5-fold CV) method to evaluate the performance of the model. This method can not only reduce over-fitting to a certain extent but also obtain as much effective information as possible from limited data [37]. More concretely, we first randomly divide the initial dataset into five sub-data sets. When the method is executed, a separate sub-data set is reserved for validating the model and the other four sub-data sets are used to train the model. This process is repeated 5 times until each sub-data set is verified once and only verified once. Finally, the average results of these 5 times are used as the performance indicators of the model. General evaluation criteria are used in this study to evaluate the performance of GCNCDA, including accuracy (Accu.), Sensitivity (Sen.), precision (Prec.), F1-Score (F1) and Matthews Correlation Coefficient (MCC). They are defined as:

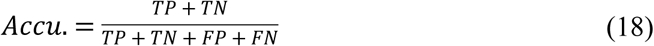

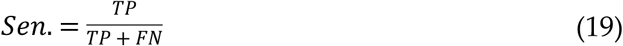

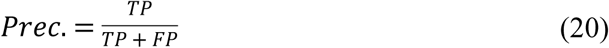

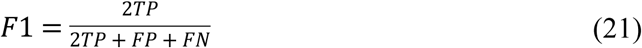

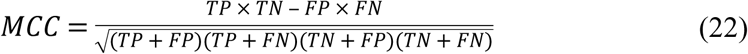

Here, TP means true positive, TN means true negative, FP means false positive, and FN means false negative. Furthermore, we also plot the Receiver Operating Characteristic (ROC) [38,39] curves of the 5-fold CV generated by GCNCDA and calculate their average area under the ROC curve (AUC) [40].

### 3.2 Model Performance Evaluation

In the experiment, GCNCDA is implemented on the benchmark dataset circR2Disease to evaluate its ability to predict potential circRNA-disease associations. The detailed results of 5-fold CV are summarized in Table 1. As can be seen from the table, GCNCDA achieved an average accuracy of 91.20% and a standard deviation of 0.74%, of which the accuracy of 5-fold experiments was 91.86%, 91.19%, 90.85%, 90.17% and 91.95%, respectively. In terms of accuracy, sensitivity, precision, F1-Score, Matthews correlation coefficient and area under ROC curve, GCNCDA obtained 92.78%, 90.03%, 91.33%, 82.55% and 90.90%, with standard deviations of 3.03%, 2.37%, 0.78%, 1.60% and 0.81%, respectively. Figure 2 plots the ROC curve generated by GCNCDA using 5-fold CV on the circR2Disease dataset. From the experimental results, we can observe that GCNCDA performs well and can effectively predict the potential disease-related circRNAs.

**Table 1.** Results of 5-fold CV generated by GCNCDA on circR2Disease dataset

| Test set | Accu. (%) | Sen. (%) | Prec. (%) | F1 (%) | MCC (%) | AUC (%) |
| --- | --- | --- | --- | --- | --- | --- |
| 1 | 91.86 | 88.97 | 94.16 | 91.49 | 83.83 | 90.93 |
| 2 | 91.19 | 93.10 | 89.40 | 91.22 | 82.45 | 90.54 |
| 3 | 90.85 | 92.52 | 89.47 | 90.97 | 81.74 | 91.24 |
| 4 | 90.17 | 91.95 | 88.96 | 90.43 | 80.38 | 89.80 |
| 5 | 91.95 | 97.39 | 88.17 | 92.55 | 84.33 | 92.00 |
| Average | **91.20±0.74** | **92.78±3.03** | **90.03±2.37** | **91.33±0.78** | **82.55±1.60** | **90.90±0.81** |

**Figure 2.**
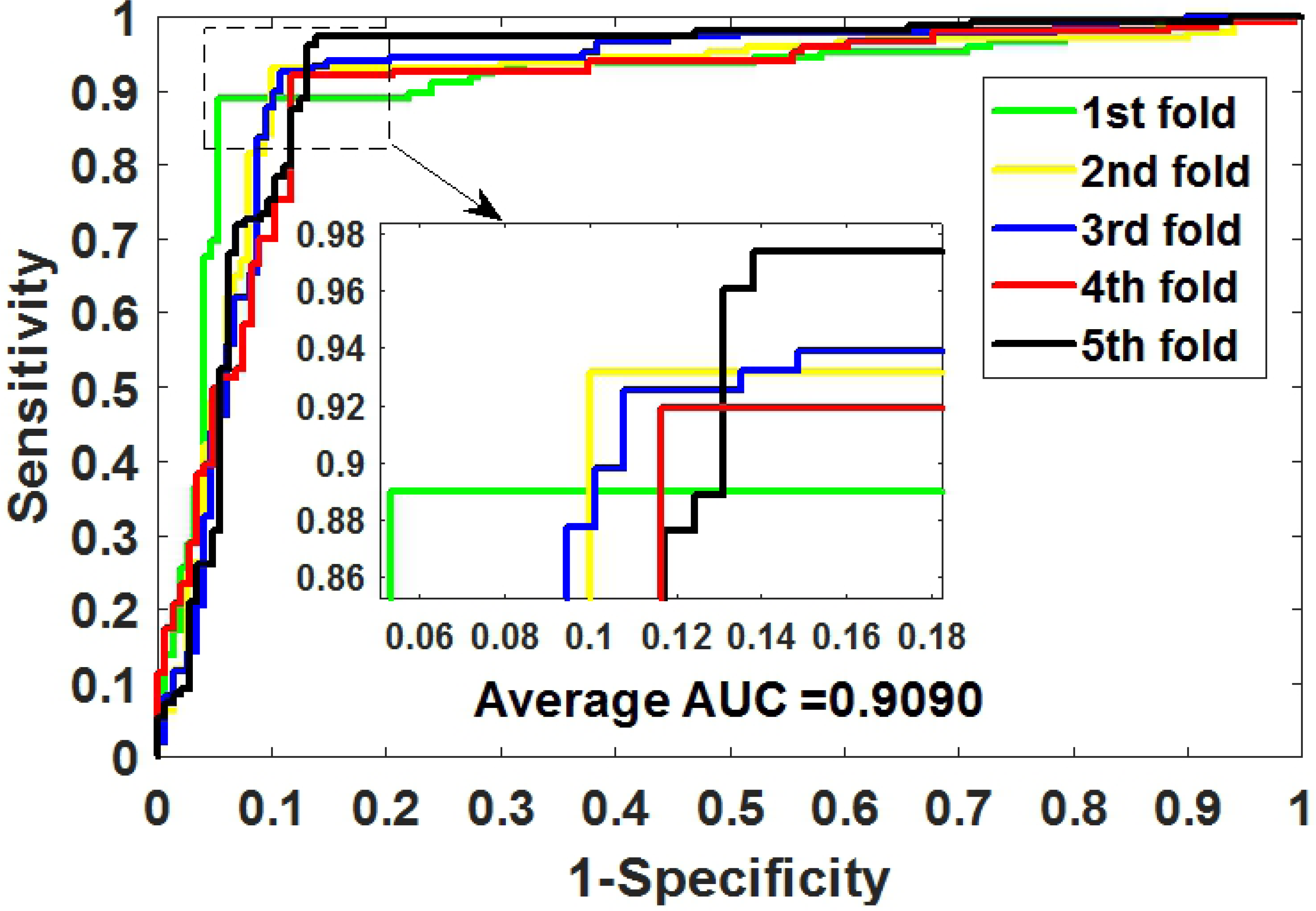
ROC curves of 5-fold CV obtained by GCNCDA on circR2Disease dataset

### 3.3 Comparison of Different Classifier Models

To evaluate the impact of the Forest PA classifier on the overall performance of GCNCDA, we compared different classifier models in this experiment. Specifically, when constructing different classifier models, we keep the other parts of the model unchanged, including the composition of descriptors and feature extraction, and only replace the Forest PA classifier with state-of-the-art Support Vector Machine (SVM) and Random Forest (RF) classifiers, respectively. The SVM model and the RF model are thus constructed and implemented on the circR2Disease dataset using 5-fold CV. Table 2 lists the results of the 5-fold CV experiments performed by these two models. Figure 3 plots the 5-fold CV ROC curves generated by the two models on the circR2Disease dataset. For the convenience of visual comparison, we display these results in the form of a histogram. As can be seen from figure 4, GCNCDA achieved the best results in accuracy, sensitivity, F1, MCC and AUC, and achieved the third result in precision, but only 2.75% lower than the best result. From the overall performance point of view, GCNCDA is better than SVM model and RF model. This result indicates that the Forest PA classifier is suitable for GCNCDA model and contributes to the improvement of the model performance.

**Table 2.** Results of 5-fold CV generated by SVM model and RF model on circR2Disease dataset

| Test set | Accu. (%) | Sen. (%) | Prec. (%) | F1 (%) | MCC (%) | AUC (%) |
| --- | --- | --- | --- | --- | --- | --- |
| 1 | 86.10 | 78.62 | 91.94 | 84.76 | 72.87 | 84.30 |
| 2 | 86.78 | 77.93 | 94.17 | 85.28 | 74.56 | 85.54 |
| 3 | 87.46 | 83.67 | 90.44 | 86.93 | 75.12 | 88.49 |
| 4 | 87.12 | 83.22 | 90.51 | 86.71 | 74.50 | 87.96 |
| 5 | 87.25 | 88.24 | 87.10 | 87.66 | 74.48 | 88.50 |
| **SVM Model** | **86.94±0.53** | **82.34±4.20** | **90.83±2.58** | **86.27±1.21** | **74.31±0.84** | **86.96±1.92** |
| 1 | 88.14 | 82.07 | 92.97 | 87.18 | 76.73 | 87.37 |
| 2 | 90.17 | 84.83 | 94.62 | 89.45 | 80.72 | 89.08 |
| 3 | 91.19 | 87.07 | 94.81 | 90.78 | 82.64 | 90.41 |
| 4 | 89.15 | 87.92 | 90.34 | 89.12 | 78.34 | 88.85 |
| 5 | 89.26 | 87.58 | 91.16 | 89.33 | 78.59 | 89.54 |
| **RF Model** | **89.58±1.15** | **85.89±2.46** | **92.78±2.01** | **89.17±1.29** | **79.41±2.30** | **89.05±1.11** |
| **GCNCDA** | **91.20±0.74** | **92.78±3.03** | **90.03±2.37** | **91.33±0.78** | **82.55±1.60** | **90.90±0.81** |

**Figure 3.**
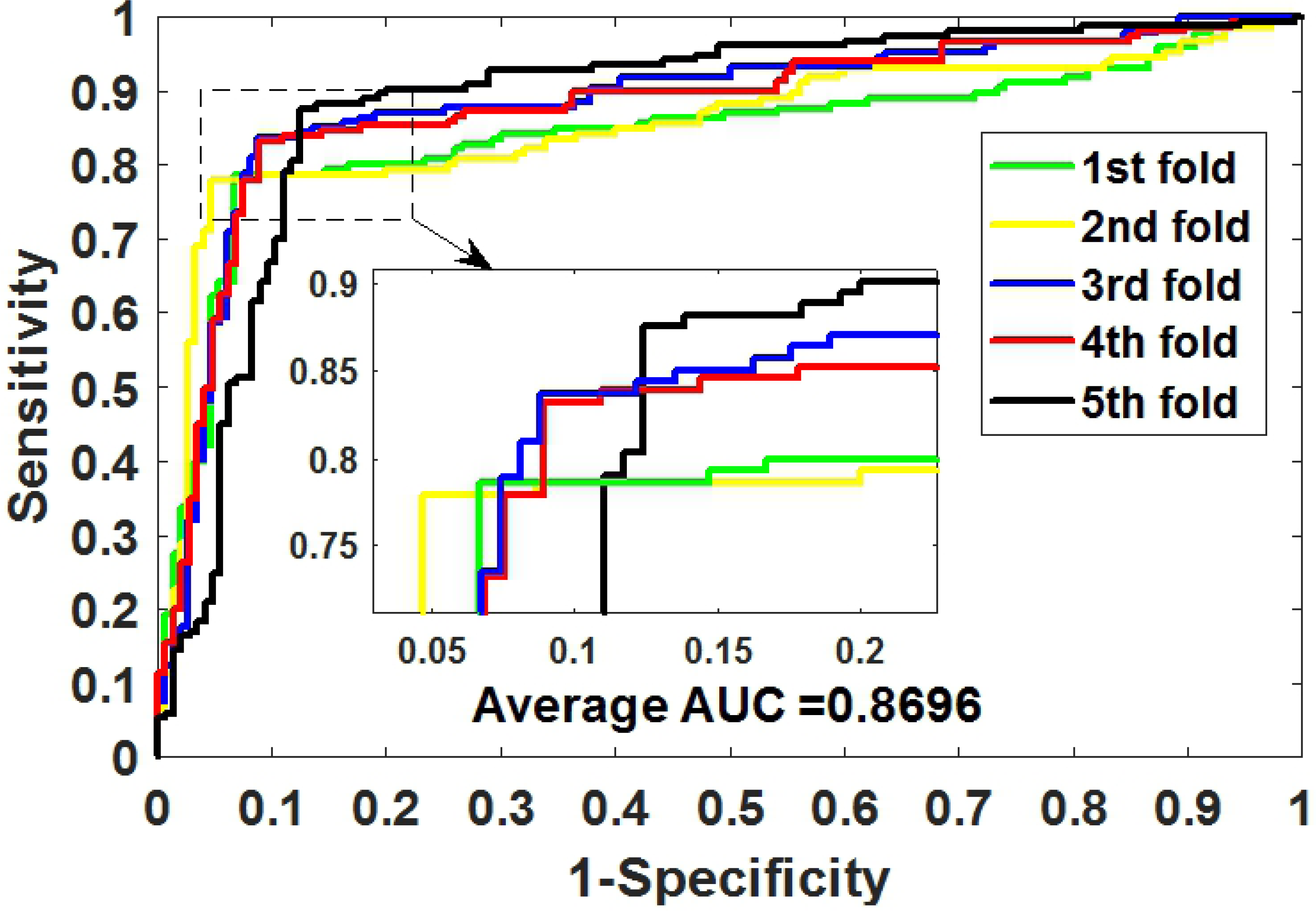
ROC curves of 5-fold CV obtained by SVM model on circR2Disease dataset

**Figure 4.**
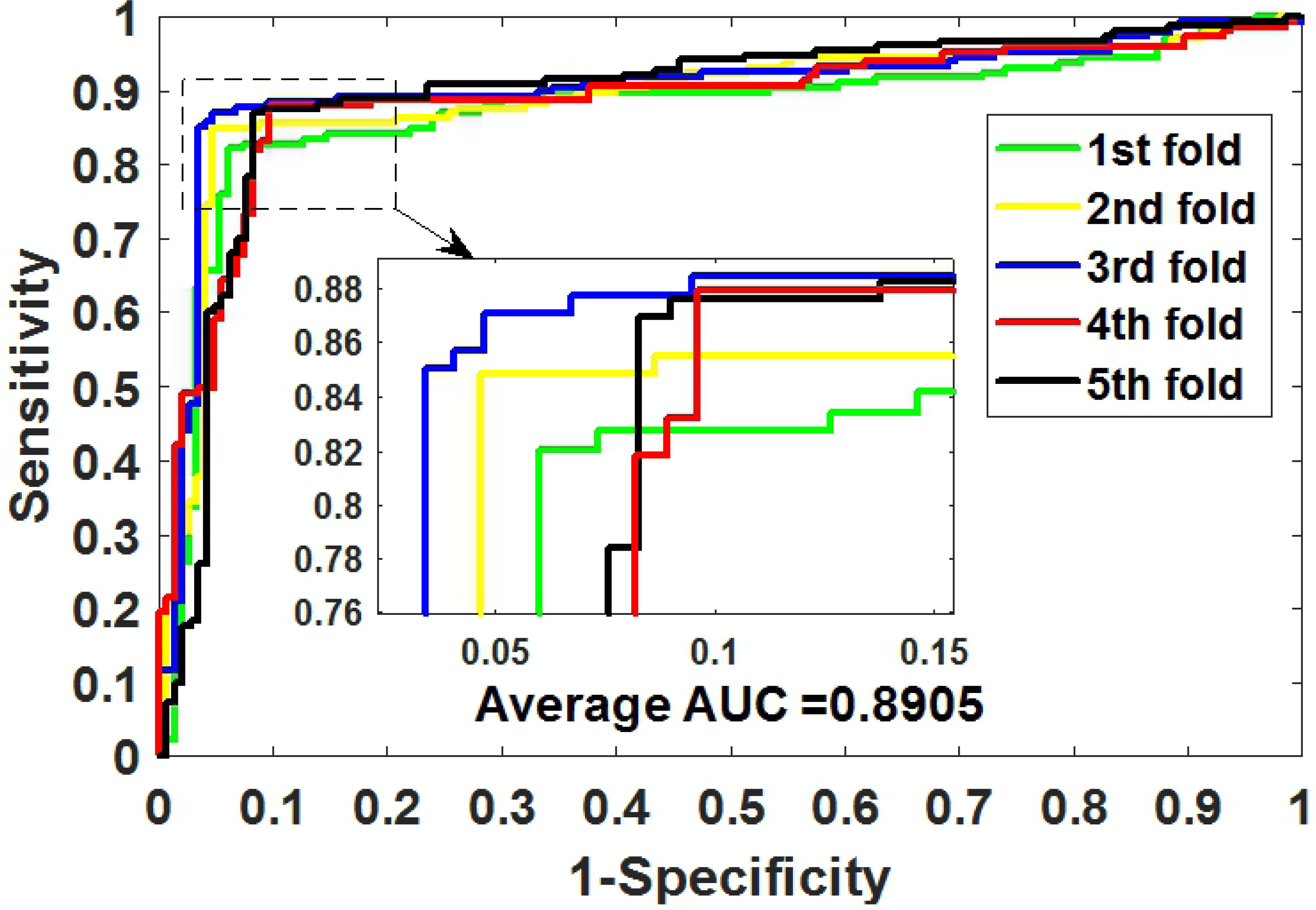
ROC curves of 5-fold CV obtained by RF model on circR2Disease dataset

### 3.4 Comparison of Different Feature Extraction Algorithms

In order to evaluate the effect of the FastGCN feature extraction algorithm on the overall performance of GCNCDA, we compared different feature extraction algorithm models in this experiment. Similar to the experiment with different classifiers, when we construct different feature extraction algorithm models, the other parts of the model are unchanged, including the composition of the descriptors and classifier. Only the Auto Covariance (AC) [41] and fast Fourier transform (FFT) [42] extraction algorithms are used instead of the FastGCN algorithm. The AC model and the FFT model are thus constructed and implemented on the circR2Disease dataset using 5-fold CV. Table 3 summarizes the results of the 5-fold CV obtained by the two models. Figure 5 plots the 5-fold CV ROC curves generated by the two models on the circR2Disease dataset. Similarly, we used a histogram to visually compare the results of the three models. As can be seen from figure 6, GCNCDA achieved the best results in all the evaluation criteria, including accuracy, sensitivity, precision, F1, MCC and AUC. The experimental results show that the FastGCN algorithm can effectively extract the advanced features of the fusion descriptor, thus helping to improve the performance of the model. In addition, from the comparison experiments of different classifiers and extraction algorithms, we can also see that the FastGCN algorithm is more helpful to the performance improvement of the model than the Forest PA classifier. This suggests that the FastGCN algorithm is the key to the GCNCDA model and plays an important role in predicting potential disease-associated circRNAs.

**Table 3.** Results of 5-fold CV generated by AC model and FFT model on circR2Disease dataset

| Test set | Accu. (%) | Sen. (%) | Prec. (%) | F1 (%) | MCC (%) | AUC (%) |
| --- | --- | --- | --- | --- | --- | --- |
| 1 | 86.44 | 93.24 | 82.14 | 87.34 | 73.55 | 85.39 |
| 2 | 85.08 | 90.67 | 81.93 | 86.08 | 70.52 | 85.83 |
| 3 | 81.36 | 86.71 | 77.50 | 81.85 | 63.23 | 81.78 |
| 4 | 85.76 | 90.13 | 83.54 | 86.71 | 71.67 | 85.74 |
| 5 | 91.95 | 97.95 | 87.20 | 92.26 | 84.54 | 93.16 |
| **ACModel** | **86.12±3.81** | **91.74±4.18** | **82.46±3.48** | **86.85±3.71** | **72.70±7.69** | **86.38±4.15** |
| 1 | 73.90 | 76.32 | 73.89 | 75.08 | 47.72 | 73.59 |
| 2 | 75.93 | 75.52 | 75.00 | 75.26 | 51.83 | 76.38 |
| 3 | 74.24 | 68.94 | 81.02 | 74.50 | 49.46 | 73.96 |
| 4 | 78.64 | 76.92 | 75.19 | 76.05 | 56.80 | 79.21 |
| 5 | 78.19 | 82.35 | 76.83 | 79.50 | 56.41 | 76.74 |
| **FFT Model** | **76.18±2.19** | **76.01±4.78** | **76.38±2.80** | **76.08±1.99** | **52.44±4.07** | **75.98±2.29** |
| **GCNCDA** | **91.20±0.74** | **92.78±3.03** | **90.03±2.37** | **91.33±0.78** | **82.55±1.60** | **90.90±0.81** |

**Figure 5.**
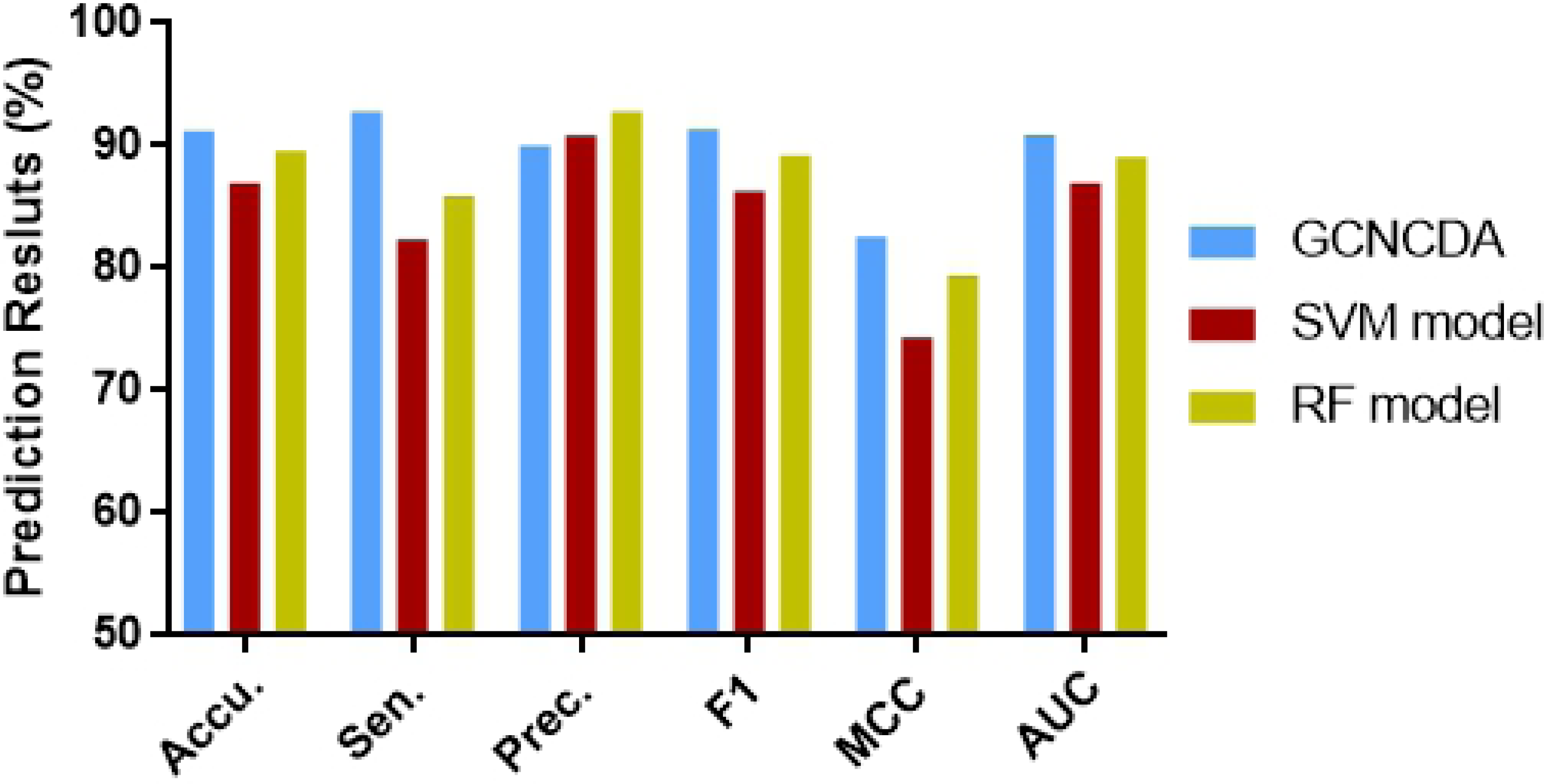
Comparison of results of different classifier models on circR2Disease dataset

**Figure 6.**
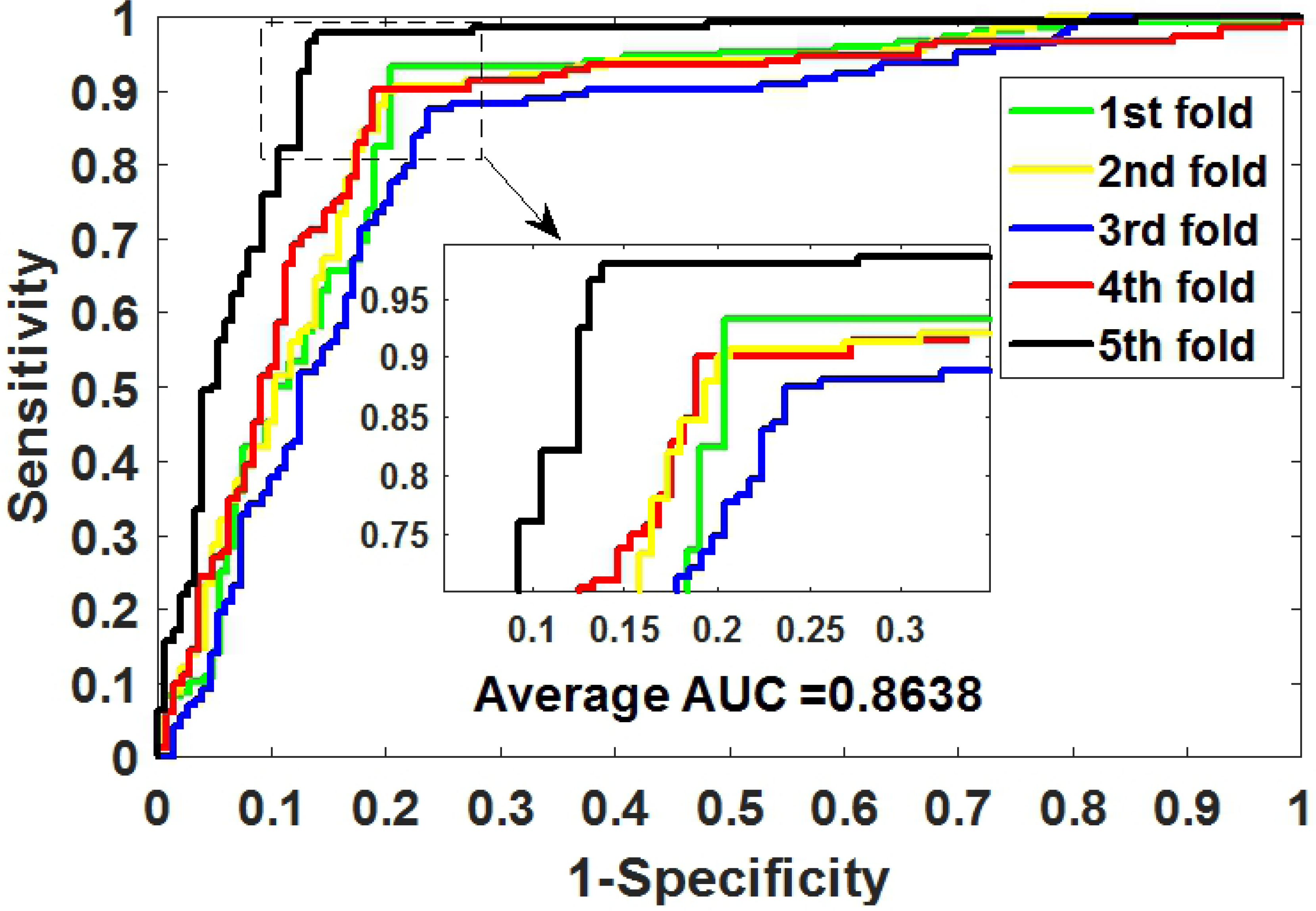
ROC curves of 5-fold CV obtained by AC model on circR2Disease dataset

**Figure 7.**
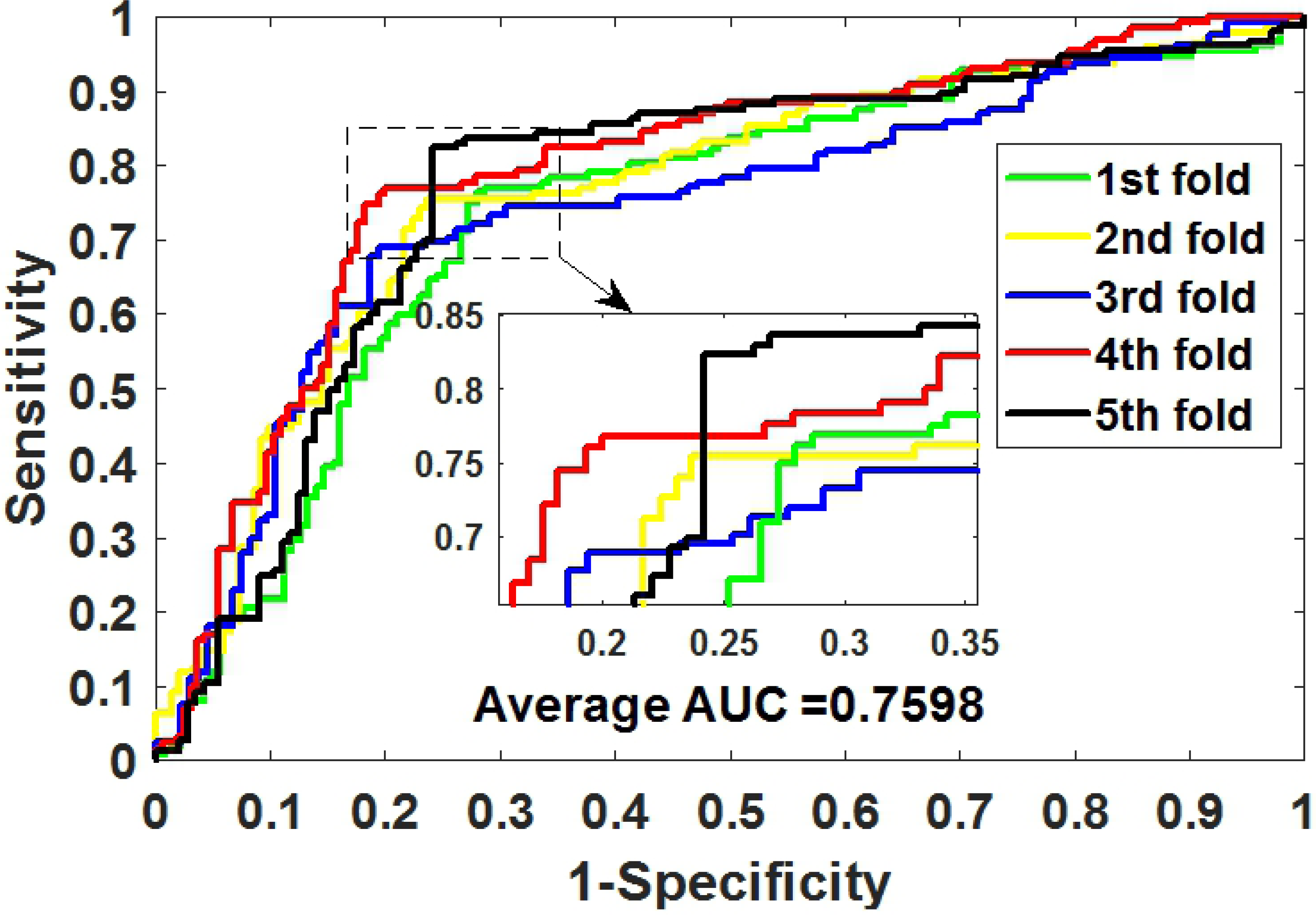
ROC curves of 5-fold CV obtained by FFT model on circR2Disease dataset

**Figure 8.**
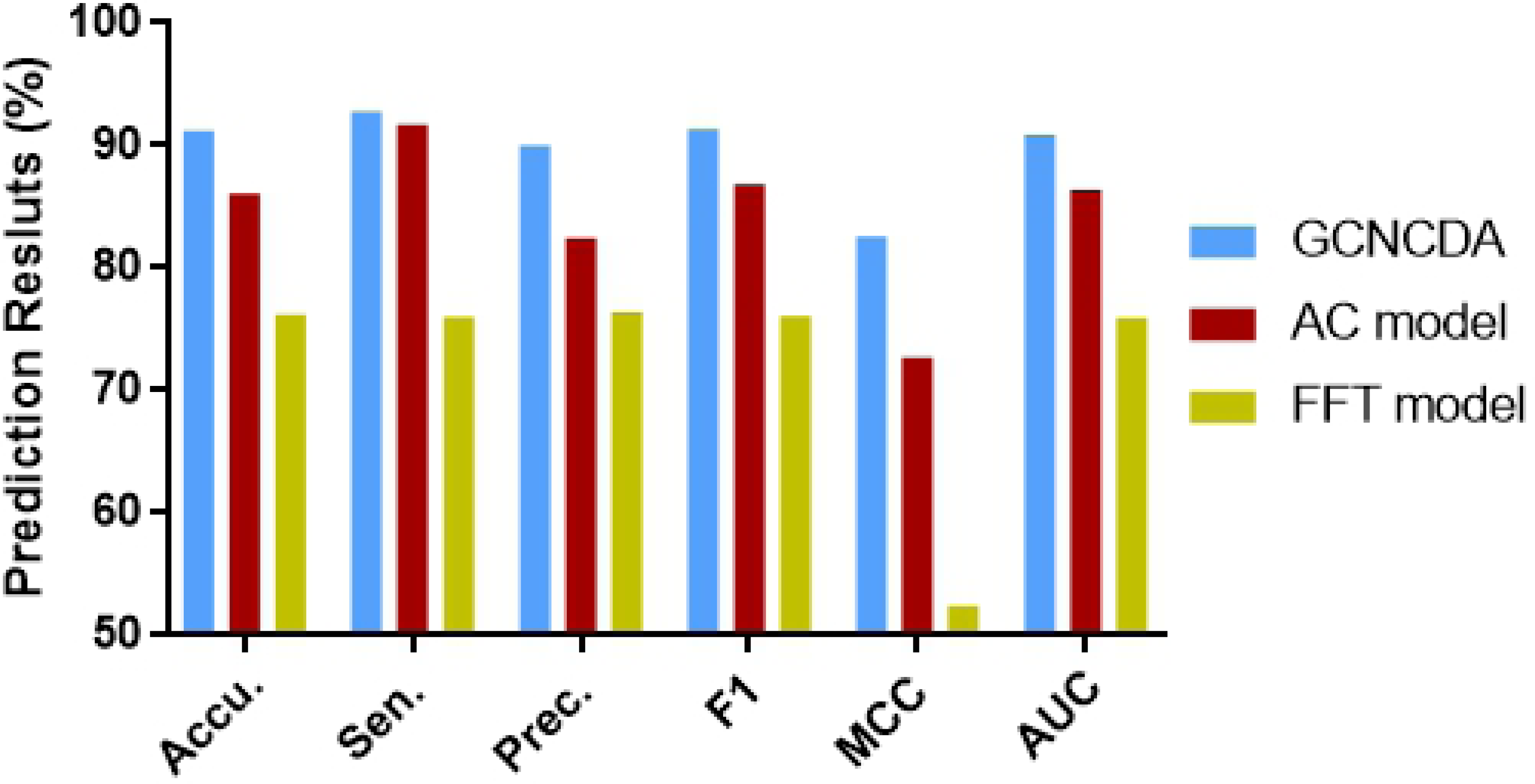
Comparison of results of different classifier models on circR2Disease dataset

### 3.5 Comparison with other existing methods

At present, some researchers have established models for predicting circRNA-disease associations based on the benchmark dataset circR2Disease, including DWNN-RLS [29], KATZHCDA [30], PWCDA [43], GHICD [43] and RWRHCD [43]. To evaluate the performance of GCNCDA, we compared it to the 5-fold CV AUC results of these models. Table 4 summarizes the 5-fold CV AUC scores generated by the various models on the same benchmark dataset circR2Disease. From the table we can see that GCNCDA is outperforms other existing methods. This indicates that the GCNCDA model, which uses the FastGCN algorithm to extract circRNA and disease fusion information features and combines the Forest PA classifier, can effectively improve the predictive performance of circRNA-disease associations.

**Table 4.** The 5-fold CV AUC scores generated by the various models on the same benchmark dataset circR2Disease

| Methods | GCNCDA | DWNN-RLS | KATZHCDA | PWCDA | GHICD | RWRHCD |
| --- | --- | --- | --- | --- | --- | --- |
| AUC | 90.90 | 88.54 | 79.36 | 89.00 | 72.90 | 66.60 |

### 3.6 Case Studies

To demonstrate the capability of GCNCDA to predict new disease-associated circRNAs based on known circRNA-disease associations, the performance of GCNCDA was further evaluated. In particular, all known circRNA-disease associations in benchmark dataset ℝ^+^ were used to train GCNCDA, and the remaining unknown circRNA-disease associations were considered candidates for testing. Then all the candidates were ranked according to the predicted score of GCNCDA. Finally, the predicted disease-circRNA associations were confirmed in the latest published literature. As a result, 10 of the top 15 circRNA-disease associations with the highest predicted scores were confirmed by relevant literature, and the detailed results are listed in table 5. It is worth noting that although some circRNA-disease associations with high predicted scores are not confirmed by existing literature, there is no denying the possibility of an association between them.

**Table 5.** Top 15 circRNA-disease associations were predicted by GCNCDA based on known circRNA-disease associations in circR2Disease database

| circRNA | Disease | Evidence (PMID) | Pub Time |
| --- | --- | --- | --- |
| hsa_circ_0004214 | cervical cancer | 28622299 | 2017 Sep |
| hsa_circ_0004214 | glioma | 28622299 | 2017 Sep |
| hsa_circ_0072088 | liver cancer | 28727484 | 2017 Jul 20 |
| hsa_circ_0013958 | lung cancer | 28685964 | 2017 Jul |
| hsa_circ_0021001 | Heart disease | unconfirmed | unconfirmed |
| circETFA | Glioma | 26873924 | 2016 May |
| hsa_circ_0070934 | Dilated cardiomyopathy | unconfirmed | unconfirmed |
| hsa_circ_0001946 | myocardial infarction | 26998750 | 2016 Mar 21 |
| hsa_circ_0001946 | glioma | 26683098 | 2016 Jan 26 |
| hsa_circ_0001955 | Immunosenescence | unconfirmed | unconfirmed |
| hsa_circ_0004458 | gastric cancer | 28544609 | 2017 Jun |
| hsa_circ_0005442 | Dilated cardiomyopathy | unconfirmed | unconfirmed |
| circRNA_010567 | Cardiovascular disease | 28412345 | 2017 Jun |
| circZFR | Bladder cancer | 27484176 | 2016 Aug |
| hsa_circ_0037911 | Heart disease | unconfirmed | unconfirmed |

## 4. Conclusion

In this study, we proposed a new computational method called GCNCDA to predict potential circRNA-disease associations. The method makes full use of the disease semantic similarity, disease and circRNA GIP kernel similarity, the known circRNA-disease association information, and extracts the high-level abstract features from them by deep learning FastGCN algorithm. The cross-validation results show that GCNCDA performs well on the benchmark dataset circR2Disease. In comparison with different classifier models, feature extraction algorithm models, and other state-of-the-art methods, GCNCDA has exhibited strong competitiveness. Furthermore, we also predicted new circRNA-disease associations based on known associations. As a result, 10 of the top 15 circRNA-disease associations with the highest predicted scores were confirmed by recently published literature. These experimental results indicate that GCNCDA is an effective method for predicting circRNA-disease associations and can provide highly reliable candidates for biological experiments. In future research, we will improve the FastGCN algorithm to help the model achieve better performance.

## Acknowledgements

This work is supported is supported in part by Awardee of the NSFC Excellent Young Scholars Program, under Grants 61722212, in part by the National Natural Science Foundation of China, under Grants 61702444, 61572506, in part by the Pioneer Hundred Talents Program of Chinese Academy of Sciences, in part by the Chinese Postdoctoral Science Foundation, under Grant 2019M653804, in part by the West Light Foundation of The Chinese Academy of Sciences, under Grant 2018-XBQNXZ-B-008. The authors would like to thank all anonymous reviewers for their constructive advices.

## Supporting Information Legends

**Supplementary Table 1.** The benchmark dataset contains 739 pairs of positive samples and 739 pairs of negative samples.

**Supplementary Table 2.** Names of 661 circRNAs involved in known circRNA-disease associations obtained from CircR2Disease database.

**Supplementary Table 3.** Names of 100 diseases involved in known circRNA-disease associations obtained from CircR2Disease database.

**Supplementary Table 4.** The MeSH dataset that provides rigorous disease classification information.

## References

1. Memczak S, Jens M, Elefsinioti A, Torti F, Krueger J, et al. (2013) Circular RNAs are a large class of animal RNAs with regulatory potency. Nature 495: 333–338.

2. Meng S, Zhou H, Feng Z, Xu Z, Tang Y, et al. (2017) CircRNA: functions and properties of a novel potential biomarker for cancer. Molecular Cancer 16: 94.

3. Jeck WR, Sharpless NE (2014) Detecting and characterizing circular RNAs. Nature Biotechnology 32: 453–461.

4. Diener T (1971) Potato spindle tuber “virus”: IV. A replicating, low molecular weight RNA. Virology 45: 411–428.

5. Hsu MT, Coca-Prados M (1979) Electron microscopic evidence for the circular form of RNA in the cytoplasm of eukaryotic cells. Nature 280: 339–340.

6. Qiu PC, Gaudette MF, Robinson DH, Crain WR (1995) Expression of the mouse testis-determining gene <em>Sry</em> in male preimplantation embryos. Molecular Reproduction & Development 40: 196.

7. Julia S, Charles G, Peter Lincoln W, Norman L, Brown PO (2012) Circular RNAs are the predominant transcript isoform from hundreds of human genes in diverse cell types. Plos One 7: e30733.

8. Hansen TB, Jensen TI, Clausen BH, Bramsen JB, Finsen B, et al. (2013) Natural RNA circles function as efficient microRNA sponges. Nature 495: 384–388.

9. Li Z, Huang C, Bao C, Chen L, Lin M, et al. (2015) Exon-intron circular RNAs regulate transcription in the nucleus. Nature structural & molecular biology 22: 256.

10. Granados-Riveron JT, Aquino-Jarquin G (2016) The complexity of the translation ability of circRNAs. BBA - Gene Regulatory Mechanisms 1859: 1245–1251.

11. Yu L, Gong X, Sun L, Zhou Q, Lu B, et al. (2016) The Circular RNA Cdr1as Act as an Oncogene in Hepatocellular Carcinoma through Targeting miR-7 Expression. Plos One 11: e0158347.

12. Tang W, Ji M, He G, Yang L, Niu Z, et al. (2017) Silencing CDR1as inhibits colorectal cancer progression through regulating microRNA-7. Oncotargets & Therapy 10: 2045.

13. Kim MK, Shin HM, Jung H, Lee E, Kim TK, et al. (2017) Comparison of pancreatic beta cells and alpha cells under hyperglycemia: Inverse coupling in pAkt-FoxO1. Diabetes Research & Clinical Practice 131: 1.

14. Floris G, Zhang L, Follesa P, Sun T (2017) Regulatory Role of Circular RNAs and Neurological Disorders. Molecular Neurobiology 54: 5156–5165.

15. Burd CE, Jeck WR, Liu Y, Sanoff HK, Wang Z, et al. (2010) Expression of Linear and Novel Circular Forms of an INK4/ARF-Associated Non-Coding RNA Correlates with Atherosclerosis Risk. Plos Genetics 6: e1001233.

16. Burd CE, Jeck WR, Yan L, Sanoff HK, Zefeng W, et al. (2010) Expression of linear and novel circular forms of an INK4/ARF-associated non-coding RNA correlates with atherosclerosis risk. Plos Genetics 6: e1001233.

17. Du WW, Yang W, Chen Y, Wu Z-K, Foster FS, et al. (2016) Foxo3 circular RNA promotes cardiac senescence by modulating multiple factors associated with stress and senescence responses. European heart journal 38: 1402–1412.

18. Lin SP, Ye S, Long Y, Fan Y, Mao HF, et al. (2016) Circular RNA expression alterations are involved in OGD/R-induced neuron injury. Biochemical & Biophysical Research Communications 471: 52–56.

19. Lukiw WJ (2013) Circular RNA (circRNA) in Alzheimer’s disease (AD). Frontiers in Genetics 4: 307.

20. Ghosal S, Das S, Sen R, Basak P, Chakrabarti J (2013) Circ2Traits: a comprehensive database for circular RNA potentially associated with disease and traits. Frontiers in genetics 4: 283–283.

21. Glažar P, Papavasileiou P, Rajewsky N (2014) circBase: a database for circular RNAs. Rna 20: 1666–1670.

22. Yang JH, Shao P, Zhou H, Chen Y-Q, Qu L-H (2010) deepBase: a database for deeply annotating and mining deep sequencing data. Nucleic Acids Research 38: D123.

23. Liu Y-C, Li J-R, Sun C-H, Andrews E, Chao R-F, et al. (2015) CircNet: a database of circular RNAs derived from transcriptome sequencing data. Nucleic acids research 44: D209–D215.

24. Fan C, Lei X, Fang Z, Jiang Q, Wu F-X (2018) CircR2Disease: a manually curated database for experimentally supported circular RNAs associated with various diseases. Database 1: 6.

25. Chen X, Han P, Zhou T, Guo X, Song X, et al. (2016) circRNADb: A comprehensive database for human circular RNAs with protein-coding annotations. Sci Rep 6: 34985.

26. Zhao Z, Wang K, Wu F, Wang W, Zhang K, et al. (2018) circRNA disease: a manually curated database of experimentally supported circRNA-disease associations. Cell death & disease 9: 475.

27. Yao D, Lei Z, Mengyue Z, Xiwei S, Yan L, et al. (2018) Circ2Disease: a manually curated database of experimentally validated circRNAs in human disease. Scientific Reports 8: 11018-.

28. Xiao Q, Luo J, Dai J (2019) Computational Prediction of Human Disease-associated circRNAs based on Manifold Regularization Learning Framework. IEEE Journal of Biomedical and Health Informatics PP: 1–1.

29. Yan C, Wang J, Wu F-X (2018) DWNN-RLS: regularized least squares method for predicting circRNA-disease associations. BMC bioinformatics 19: 520.

30. Fan C, Lei X, Wu F-X (2018) Prediction of CircRNA-Disease Associations Using KATZ Model Based on Heterogeneous Networks. International journal of biological sciences 14: 1950.

31. Macintyre G, Jimeno YA, Ong CS, Verspoor K (2014) Associating disease-related genetic variants in intergenic regions to the genes they impact. Peerj 2: e639.

32. Wang L, You Z-H, Chen X, Li Y-M, Dong Y-N, et al. (2019) LMTRDA: Using logistic model tree to predict MiRNA-disease associations by fusing multi-source information of sequences and similarities. PLoS computational biology 15: e1006865.

33. Xiang Z, Qin T, Qin ZS, He Y (2013) A genome-wide MeSH-based literature mining system predicts implicit gene-to-gene relationships and networks. BMC systems biology 7: S9.

34. Wang D, Wang J, Lu M, Song F, Cui Q (2010) Inferring the human microRNA functional similarity and functional network based on microRNA-associated diseases. Bioinformatics 26: 1644–1650.

35. Jie C, Ma T, Cao X (2018) FastGCN: Fast Learning with Graph Convolutional Networks via Importance Sampling. International Conference on Learning Representations.

36. Adnan MN, Islam MZ (2017) Forest PA: Constructing a decision forest by penalizing attributes used in previous trees. Expert Systems with Applications 89: 389–403.

37. Wang L, You Z-H, Yan X, Xia S-X, Liu F, et al. (2018) Using Two-dimensional Principal Component Analysis and Rotation Forest for Prediction of Protein-Protein Interactions. Scientific reports 8: 12874.

38. Zweig MH, Campbell G (1993) Receiver-operating characteristic (ROC) plots: a fundamental evaluation tool in clinical medicine. Clinical chemistry 39: 561–577.

39. Swets JA (1988) Measuring the accuracy of diagnostic systems. Science 240: 1285.

40. Bradley AP (1997) The use of the area under the ROC curve in the evaluation of machine learning algorithms. Pattern recognition 30: 1145–1159.

41. Guo Y, Yu L, Wen Z, Li M (2008) Using support vector machine combined with auto covariance to predict proteinprotein interactions from protein sequences. Nucleic Acids Research 36: 3025–3030.

42. Lin S-J, Alnaffouri T, Han Y, Chung W-H (2016) Novel Polynomial Basis with Fast Fourier Transform and Its Application to Reed-Solomon Erasure Codes. IEEE Transactions on Information Theory 62: 6284–6299.

43. Lei X, Fang Z, Chen L, Wu F-X (2018) PWCDA: Path Weighted Method for Predicting circRNA-Disease Associations. International journal of molecular sciences 19: 3410.

